# Comparison of methods for milk pre-processing, exosome isolation, and RNA extraction in bovine and human milk

**DOI:** 10.1101/2020.08.14.251629

**Authors:** Sanoji Wijenayake, Shafinaz Eisha, Zoya Tawhidi, Michael A. Pitino, Michael A. Steele, Alison S. Fleming, Patrick O. McGowan

**Affiliations:** Center for Environmental Epigenetics and Development, Department of Biological Sciences, University of Toronto Scarborough, Toronto. ON, Canada; Department of Cell and Systems Biology, University of Toronto, Toronto, ON, Canada; Department of Psychology, University of Toronto, Toronto, ON, Canada; Department of Physiology, University of Toronto, Toronto, ON, Canada; Department of Nutritional Sciences, University of Toronto, Toronto, ON, Canada; Translational Medicine Program, The Hospital for Sick Children, Toronto, ON, Canada; Department of Animal and Poultry Science, University of Guelph, Guelph, ON, Canada; Department of Psychology, University of Toronto, Mississauga, Mississauga, ON, Canada

**Keywords:** Milk exosomes, bioactive compounds, extracellular vesicles, ExoQuick solution, ultracentrifugation, human, bovine

## Abstract

Milk is a highly complex, heterogeneous biological fluid that contains bioactive, membrane-bound extracellular vesicles called exosomes. Characterization of milk-derived exosomes (MDEs) is challenging due to the lack of standardized methods that are currently being used for milk pre-processing, exosome isolation, and RNA extraction. In this study, we tested: 1) three pre-processing methods to remove cream, fat, and casein proteins from bovine milk to determine whether pre-processing of whole milk, prior to long-term storage, improves MDE isolations, 2) two commonly-used exosome isolation methods, and 3) four extraction protocols for obtaining high quality MDE RNA from bovine and human milk. MDEs were characterized via Transmission Electron Microscopy (TEM) and Nanoparticle Tracking Analysis (NTA). We also present an optimized method of TEM sample preparation and isolation of total soluble protein from MDEs. Our results indicated that: 1) pre-processing of bovine milk prior to storage does not affect the final exosome yield or the purity, 2) ExoQuick precipitation is better suited for MDE isolation than ultracentrifugation for bovine and human milk, and 3) TRIzol LS produced the highest RNA yield in bovine milk, whereas TRIzol LS, TRIzol+RNA Clean and Concentrator, and TRIzol LS+RNA Clean and Concentrator methods can be used for human milk.

## Introduction

Maternal milk is the primary nutritional source of newborn mammals. Unlike standardized infant formula, which has a narrow macro/micronutrient range, maternal milk composition is highly dynamic and varies across lactational stages, the circadian cycle, maternal age, ethnicity, diet, and gestational age (1–3). Mammalian milk is a highly complex and heterogeneous solution that contains protein, lipids, carbohydrates, minerals, vitamins, active enzymes, hormones, immune factors, and microbiota (1,2,4,5). Mammalian milk is also biologically customized to fit the physiological, neurodevelopmental, and immune requirements of offspring as they age (1,2,6,7). In particular, human colostrum, is a rich source of immunological components including IgA, lactoferrin, leukocytes, and human milk oligosaccharides (HMOs), and is a vital source of early-life immune programming. In contrast, transition and mature milk are mainly tailored to meet the nutrient and energy demands of the growing offspring (8,9).

Recently, maternal milk was found to contain functional microRNAs (miRNAs) encapsulated in protective milk-derived exosomes (MDEs) (10–18). MDEs (a subtype of extracellular vesicles (EVs)) range from 30-150 nm in size and were identified in many mammals including humans, cows, rodents, goats, pigs, and marsupials (17,19,20). MDEs are secreted from mammary gland epithelial cells (MECs), can travel across offspring’s intestinal endothelium post-ingestion into circulation, and are taken up by surrounding tissues (14). Upon intake into cells, MDEs may release their cargo and regulate cellular functions of the recipient cells (10–14,16,17,20–22). Lactation-specific miRNAs have been shown to induce post-transcriptional regulation of target mRNA in various recipient tissues (11,18,20,21,23). MDEs have also been shown to cross biological barriers in vitro and in vivo (14,24), increasing interest in their potential roles in nutritional and agricultural research, translational medicine, and drug therapeutics (25–27).

Thus, the isolation and characterization of MDEs have become an important area of research, and there is a need to establish standardized and reproducible methods to isolate and quantify MDEs. In 2014, the International Society for Extracellular Vesicles (ISEV) published a Position Editorial detailing the minimal requirements and recommendations for the identification and characterization of extracellular vesicles and their proposed functions (28). Although these guidelines are regularly updated, they indicate that isolation of intact MDEs with high purity and yield remains a challenge largely due to the numerous handling, processing, and isolation techniques currently used in EV research. The use of non-standardized techniques is more of an issue in MDE research due to the high intra- and interspecific variability that naturally exist across milk samples. In particular, some studies report that changes in temperature and long-term storage of maternal milk do not affect milk composition, integrity, and the final yield of isolated MDEs, while other studies have found that the recovery of MDEs from maternal milk is influenced by sample collection and pre-processing steps (12,29–31). Consequently, here we tested three pre-processing techniques on unpasteurized, whole bovine milk to determine whether removing cream, fat globules and/or casein proteins prior to ultracold storage is required to obtain high quality MDEs.

The purity of the isolated exosomes can vary due to the presence of contaminating particles, other EVs, viscosity of the sample, the presence of milk proteins, and nucleic acids that are often precipitated alongside MDEs (32–34). Ultracentrifugation-based (UC) isolations and polymer-based precipitation techniques (i.e. ExoQuick (EQ) reagent) are two of the more commonly used exosome isolation methods (10,11,36–41,12–14,26,32–35). Differential centrifugation combined with UC is considered the gold standard for exosome isolation, although ultra-high speeds and continuous handling of the samples can result in degradation and low recovery rates. UC is also time consuming, requires specialized equipment and a large starting volume (26,35). In comparison, EQ precipitation has a high recovery rate, is a faster method, does not require a large sample volume (13,26,32,33,35,37,38), and can produce higher miRNA yield with greater purity than other techniques (42). However, EQ precipitation may also co-precipitate contaminants and other EVs that can interfere with downstream applications (26,32,33). Therefore, further studies are necessary to confirm the suitability of both isolation techniques in MDE research across species, especially taking into account the downstream applications, including RNA and protein yield resulting from each method. As such, we tested two exosome isolation methods: EQ precipitation and differential UC using unpasteurized bovine and human milk. Transmission electron microscopy (TEM) and Nanoparticle Tracking Analysis (NTA) were used to characterize the isolated exosomes.

Since the first identification of RNA in exosomes in 2007 (25), numerous extraction methods have been used for RNA profiling via real time quantitative PCR (RT-qPCR), microarrays, and RNA sequencing. However, enrichment and molecular profiling of MDEs remains technically challenging (37) due to the variability in RNA extraction protocols and commercially available RNA extraction kits that are often utilized in EV research. Here, we tested four MDE-based RNA extraction protocols: three commercially available kits (QIAzol + miRNeasy Mini Kit, TRIzol + RNA Clean and Concentrator Kit (RCC), TRIzol LS + RCC) and an inhouse phenol-based extraction method (TRIzol LS) (43) to identify the most reproducible and optimal RNA extraction method that can be used to generate high quality RNA (≥17 nucleotides) from bovine and human MDEs.

The overall aim of the current study was to compare methods of milk pre-processing, MDE isolation techniques, and RNA extraction protocols that are currently being used in the EV field to identify the most robust and reproducible techniques that can be used to standardize MDE research. Our results may be useful for the selection of purification methods in future studies using human and/or bovine milk where sample volumes are limited, samples are subjected to extended storage times, and are frozen upon collection (e.g. human donor milk banks).

## Materials and Methods

### Bovine milk collection and processing

Unpasteurized bovine milk was obtained from Loa-De-Mede Holsteins Farm (Oshawa, Ontario, Canada) from 3 different dairy cows. 100 mL of bovine milk per dairy cow was collected into sterile, DNA/RNase-free conical tubes via hand milking, stored at 4 °C and transported to the University of Toronto, Scarborough for analysis. All bovine samples were pooled to remove variability in milk composition across dairy cows but, processed separately to ensure independent sample extractions.

Three different processing steps were conducted (n=3 independent trials/group) to test whether removal of cream, fat globules, and casein proteins prior to long-term storage at −80 °C may impact MDE isolation and characterization efficiency (Supplementary Fig. 1): Group (G) 1) unprocessed, whole milk stored at −80 °C, where samples were immediately frozen at −80 °C upon arrival; G2) processed milk without fat globules and cream, where bovine milk was centrifuged twice at 3,000 x g for 10 min at room temperature (RT) and the supernatant was collected and stored at −80 °C; and G3) isolated whey fraction without fat globules, cream, cellular debris, and casein proteins. G3 bovine milk samples were processed as per G2 procedure plus 2x centrifugations at 1,200 x g for 10 min at 4 °C to remove residual fat globules and cellular debris. Subsequently, the defatted supernatants were centrifuged 2x at 21,500 x g for 30 min at 4 °C followed by a subsequent centrifugation at 21,500 x g for 1 h to pellet casein proteins. The supernatants were filtered once through 0.45 μM (FroggaBio; SF0.45PES) and 0.22 μM (FroggaBio; SF0.22PES) PES syringe filters to remove residual cell debris. Isolated whey portion of bovine milk was stored at −80 °C for later use.

### Human milk collection

Expressed human milk from 2 different donors (500mL/donor) was obtained from the Roger Hixon Ontario Human Milk Bank (Toronto, Ontario, Canada). The unpasteurized samples of human milk used in this study contained a bacterial load >5 × 10^7^ colony forming units/L and were therefore not suitable for processing or dispensing for human consumption as per the policies of the milk bank. 500 mL of human milk/donor were collected into sterile collection bags, frozen at −20 °C immediately upon collection, remained frozen during transport to the milk bank, and subsequently stored at −20 °C till use. Next, the milk samples were thawed overnight at 4 °C and were pooled but processed separately to ensure independent sample extractions. Note: the samples were used in the analysis within 8 months of storage.

### Exosome Isolation

Two main isolation methods that are frequently used in exosome-isolation and characterization studies, including EQ precipitation and UC, were compared to determine the most efficient method for isolating MDEs from the whey portion of bovine and human milk (n=3 independent isolations/method) (Supplementary Fig. 2).

### Method 1: ExoQuick Precipitation

Human and G1 bovine milk samples were centrifuged at 2,000 x g for 10 min at 4 °C to remove upper cream layer. The supernatants were collected carefully and centrifuged again at 12,000 x g for 30 min at 4 °C to remove fat cells and globules. Finally, the supernatants were further centrifuged at 12,000 x g for 5 min at 4 °C to pellet cell debris. Supernatants were isolated and filtered once through 0.45 μM syringe filters to eliminate traces of cellular debris. G2 bovine samples were filtered once through 0.45 μM (FroggaBio; SF0.45PES) syringe filters to eliminate cellular debris. G3 whey fractions were used directly in the exosome precipitation reaction.

MDEs were isolated from G1-G3 bovine and human milk samples (1.5 mL/sample) using the EQ reagent (System Biosciences: EXOQ5A-1) (13,14,38,40,44). EQ reagent was added to all samples (1: 0.2, v/v), mixed by inversion, and incubated for 12 h at 4 °C to enhance precipitation. Post incubation, all samples were centrifuged at 18,000 x g for 45 min at 4 °C to pellet the exosomes. The pelleted exosomes were re-suspended in 200-400 μL of 1x-filtered PBS (determined based on the size of the pellet). The supernatants were used as the negative control for subsequent experiments.

### Method 2: Ultracentrifugation

An optimized version of the UC method as previously described (10,11,35,39,40,45) was used. Human and G1 bovine milk samples were centrifuged twice at 3,000 x g for 10 min at RT to remove the upper cream layer. The supernatants were centrifuged twice at 1,200 x g for 10 min at 4 °C, followed by two top-speed centrifugations at 21,500 x g for 30 min at 4 °C and a final centrifugation at 21,500 x g for 1 h at 4 °C to remove fat globules and casein proteins. The supernatants were filtered through 0.45 μM and 0.22 μM PES syringe filters to remove cell debris and residual fat cells. G2 bovine samples were centrifuged twice at 1,200 x g for 10 min at 4 °C, followed by two top-speed, centrifugations at 21,500 x g for 30 min at 4 °C and a final centrifugation at 21,500 x g for 1 h at 4 °C to remove casein proteins. Similar to G1 bovine samples, the supernatants were filtered through 0.45 μM and 0.22 μM PES syringe filters to remove cell debris and residual fat cells. G3 bovine samples were directly used in the ultracentrifugation step. All whey fractions were centrifuged at 100,000 x g for 90 min using a SW55 TI swing bucket ultracentrifuge at 4 °C. The pellets were re-suspended in 200 μL of filtered 1X PBS. The supernatants were used as the negative control for all subsequent experiments.

### Exosome Visualization

The isolated MDEs were visualized by TEM with negative staining using an optimized sample preparation technique. Four-hundred mesh carbon-coated copper grids (Electron Microscopy Sciences; CF400-CU-50) were incubated for 5 min with 10 μL of isolated MDEs. Three consecutive wash steps with 20 μL of ddH_2_O (2 min each) were done to minimize crystallization and coagulation of milk residue. All copper grids were negatively stained with 20 μL of 2 % uranyl acetate for 5 min at RT (Supplementary Fig. 3). All excess reagents were removed with filter paper to ensure a 100 nm thickness and all grids were dried under an incandescent light for 2 min. The copper grids were observed and photographed using a Hitachi H-7500 transmission electron microscope with a Megaview III camera (Olympus).

### Exosome Quantification

Particle size and concentration of isolated exosomes and negative controls were quantified using Nanoparticle Tracking Analysis (Malvern Instruments Ltd.; NanoSight NS300) as per manufacturer’s instructions at the Structural and Biophysical Core Facility located in the Hospital for Sick Children (Toronto, Ontario, Canada). NTA utilize the properties of Brownian motion and light scattering to measure particle size and concentration (particles/mL) of EVs. The software tracks individual particles frame by frame and calculates particle size based on Stokes-Einstein equation (46). A 1:700 dilution factor for bovine samples and a 1:300 dilution factor for human samples were used for the analysis. Standard curves ranging from 1:100 to 1:700 (v/v in 1X PBS) were run per species to determine the correct dilution range (60-100 particles/frame). An absolute control of 1X PBS was also assessed. Processing settings consisted of a detection threshold of 8, camera level of 15, and 3 replicates of 30 s captures. The laser type was Blue 488 nm.

### RNA Extraction of MDEs

Four RNA extraction methods that are commonly used in exosome studies (7–12,18,20,27,32,34,35,37,39,41,43–46) were compared to identify the most repeatable and suitable method to obtain high quality MDEs from bovine and human milk (n=6 independent extractions/protocol) (Supplementary Fig. 4).

### Method 1: QIAzol + miRNeasy Mini Kit

QIAzol lysis reagent combined with miRNeasy Mini Kit (Qiagen; 217004) was used as per the manufacturer’s instructions with slight modifications. QIAzol reagent was added to 200 μL of isolated milk exosomes (5:1, v/v). All samples were homogenized by pipetting 20 x followed by aspirating 20 x with 18-gauge needles and incubated for 5 min at RT. Chloroform was added to each sample (1:1, v/v to the starting sample), then all samples were shaken vigorously for 15 s to mix and incubated for 3 min at RT. Phase separation was done by centrifuging at 12,000 x g for 15 min at 4 °C. The upper aqueous phase was collected for RNA extraction and 100 % ethanol (1.5:1, v/v) was added and pipetted to mix. The lower phenol layer was kept aside for protein extractions. All content, including any precipitate, was transferred to RNeasy mini columns and centrifuged at 10,000 x g for 15 s at RT. Columns were washed according to the manufacturer’s instructions. Post washing, columns were centrifuged at the maximum speed for 5 min to remove ethanol contamination. RNA was eluted with 50 μL DNA/RNase free ddH_2_O. Columns were incubated for 10 min after adding DNA/RNase-free water and re-eluted to increase RNA yield. RNA concentration (ng/μL) and quality (A260/A280; A260/A230) were determined using a Nanodrop Spectrophotometer (Thermo Scientific; ND-2000C).

### Method 2: TRIzol LS

TRIzol LS reagent (Thermo Scientific; 10296010) was used as described in (43) with minor modifications. TRIzol LS reagent was added to 200 μL of isolated milk exosomes (3:1, v/v). All samples were homogenized by pipetting 20 x followed by aspirating 20 x with 18-gauge needles and incubated for 5 min at RT. 200 μL of chloroform was added to each sample. Samples were mixed by shaking for 30 s and incubated for 10 min at RT. Phase separation was done by centrifuging samples at 12,000 x g for 15 min at 4 °C. The upper aqueous phase was collected for RNA extraction, while the lower phenol phase was kept aside for protein isolation. 10 % sodium acetate (3M, pH 5.5), 4 μL of glycogen, and 100 % ethanol (2.5:1, v/v) of the volume of aqueous phase were added per sample. Samples were mixed and incubated overnight at −80 °C to facilitate RNA precipitation. Post incubation, samples were centrifuged at 16,000 x g for 30 min at 4 °C to pellet the RNA. Subsequently, RNA pellets were washed with 500 μL of 70 % ethanol and centrifuged at 16,000 x g for 5 min at 4 °C. Ethanol was aspirated and pellet was centrifuged again at top speed for 1 min to remove any residual ethanol. Of note, that this step was extremely important for the proper removal of ethanol contamination. The pellet was air-dried for 10 min and re-suspended in 32 μL of RNase-free ddH_2_O. RNA concentration (ng/μL) and quality (A260/A280; A260/A230) were determined using a Nanodrop Spectrophotometer (Thermo Scientific; ND-2000C).

### Method 3: TRIzol + RNA Clean and Concentrator Kit

MDEs were lysed using TRIzol reagent (Thermo Scientific; 15596026) as per the manufacturer’s instructions with modifications. Cold TRIzol reagent was added to 200 μL of isolated MDEs (5:1, v/v). Samples were homogenized by pipetting 20 x followed by aspirating 20 x with 18-gauge needles and incubated for 5 min at RT. Subsequently, 200 μL of chloroform was added to the samples (0.2:1, v/v to TRIzol), vortexed for 30 s, and incubated for 3 min at RT. Samples were centrifuged at 12,000 x g for 15 min at 4 °C to induce phase separation. The colorless, upper aqueous phase, containing total soluble RNA, was collected while the lower phenol phase was kept aside for protein isolation. Following TRIzol phase separation, 100 % ethanol was added to the aqueous phase (1:1, v/v) and transferred to RNA Clean and Concentrator (RCC) ™ −5 kit (Zymo Research; R1013) and centrifuged at 16,000 x g for 30 s. RCC kit can be used to isolate ultra-pure, total RNA (≥17 ntd in length). Subsequently, the columns were washed once with RNA prep buffer and twice with RNA wash buffer (supplied with the kit) as per the manufacturer’s instructions. After the last wash, the columns were centrifuged on max speed for 5 min to remove residual ethanol. RNA was eluted with 40 μL of DNA/RNase-free water. Note: columns were incubated for 10 min after adding DNA/RNase-free water and re-eluted to increase RNA yield. RNA concentration (ng/μL) and quality (A260/A280; A260/A230) were determined using a Nanodrop Spectrophotometer (Thermo Scientific; ND-2000C).

### Method 4: TRIzol LS + RNA Clean and Concentrator Kit

MDEs were lysed using TRIzol LS reagent (Thermo Scientific; 10296010) as per the manufacturer’s instructions with modifications. Cold TRIzol LS reagent was added to 200 μL of isolated milk exosomes (3:1, v/v). Samples were homogenized by pipetting 20 x followed by aspirating 20 x with 18-gauge needles and incubated for 5 min at RT. Subsequently, 200 μL of chloroform was added to the samples (0.3:1, v/v to TRIzol LS), vortexed for 30 s and incubated for 3 min at RT. Samples were centrifuged at 12,000 x g for 15 min at 4 °C to induce phase separation. The colorless, upper aqueous phase, containing total soluble RNA was collected while the lower phenol phase was kept aside for protein isolation. All the steps involving the use of RCC kit are identical to that of TRIzol +RCC method. RNA concentration (ng/μL) and quality (A260/A280; A260/A230) were determined using a Nanodrop Spectrophotometer (Thermo Scientific; ND-2000C).

All RNA samples were separated via gel electrophoresis (1 % TAE agarose, w/v) for 60 min at 300 mV to visualize RNA integrity and traces of cellular RNA contamination of the isolated exosome fractions. 1 kB DNA ladder (250 bp – 4,000 bp) (Genedirex; DM101-R500) and a cellular RNA control were run alongside the samples.

### Protein Isolation

Total soluble protein was isolated from the lower phenol phase from all four RNA extraction protocols listed above, as per manufacturer’s instructions with modifications. 100 % ethanol was added (1:0.3, v/v) to the samples, incubated for 3 min at RT, and centrifuged at 2,000 x g for 5 min at 4 °C to pellet gDNA. Isopropanol (1:1.5, v/v) was added to the resulting supernatant and incubated for 10 min at RT. Subsequently, the samples were centrifuged at 12,000 x g for 10 min at 4 °C to pellet the proteins. The pellets were washed with 0.3 M guanidine hydrochloride in 95 % ethanol (1:2, v/v) and incubated for 20 min at RT. The washing step was repeated twice more. Finally, the pellets were washed with 2 mL of 100 % ethanol and incubated for another 20 min at RT. All centrifugation steps were done at 7,500 x g for 5 min at 4 °C. The pellets were air-dried for 10 min to remove residual ethanol and phenol contamination. The pellets were re-suspended in 200 μL of 1 % SDS and incubated in a water bath at 50 °C for 20 min. To enhance the solubility of proteins, the samples were incubated for 12 h at 4 °C and centrifuged at 10,000 x g for 10 min at 4 °C to remove insoluble material. Protein concentration was determined using a Pierce ™ Bicinchoninic Acid (BCA) Protein Microplate Assay (Thermo Scientific; 23225) and BSA standards ranging from 2000 μg/mL to 0 μg/mL.

### Statistical Analysis

Statistical analysis was conducted using SPSS statistical software (IBM Corp.), and figures were created using GraphPad Prism Version 7 and BioRender.com. A Shapiro-Wilk test was used to assess normality. The data were normally distributed (p>0.05) and as such parametric analyses were carried out. One-way analysis of variance (ANOVA) was used to test for main effect of bovine milk pre-processing (G1-G3). Three-way ANOVA was used to test for main effect of exosome fractionation (pellet and supernatant), exosome isolation methods (EQ and UC), RNA extraction protocols, and their interactions. Tukey post-hoc analysis was used to conduct all pairwise comparisons. Relationships were considered statistically significant at p ≤ 0.05.

## Results

### Pre-processing of Bovine Milk

NTA (Fig 1A-B), RNA (Fig. 1C-E), and protein (Fig. 1F) results indicated no significant effect of pre-processing on bovine milk samples (NTA: main effect of pre-processing, (*F*_(2, 23)_ = 0.440, *p* = 0.654); RNA concentration: (*F*_(2, 95)_ = 1.050, *p* = 0.354); RNA purity–A260/A280: (*F*_(2, 95)_ = 1.313, *p* = 0.274); A260/A230: (*F*_(2, 95)_ = 0.568, *p* = 0.569); protein concentration: (*F*_(2,95)_ = 0.431, *p* = 0.651). However, slight differences in exosome quality were visually observed across G1-G3 via TEM (Fig. 2–3).

**Figure 1.**
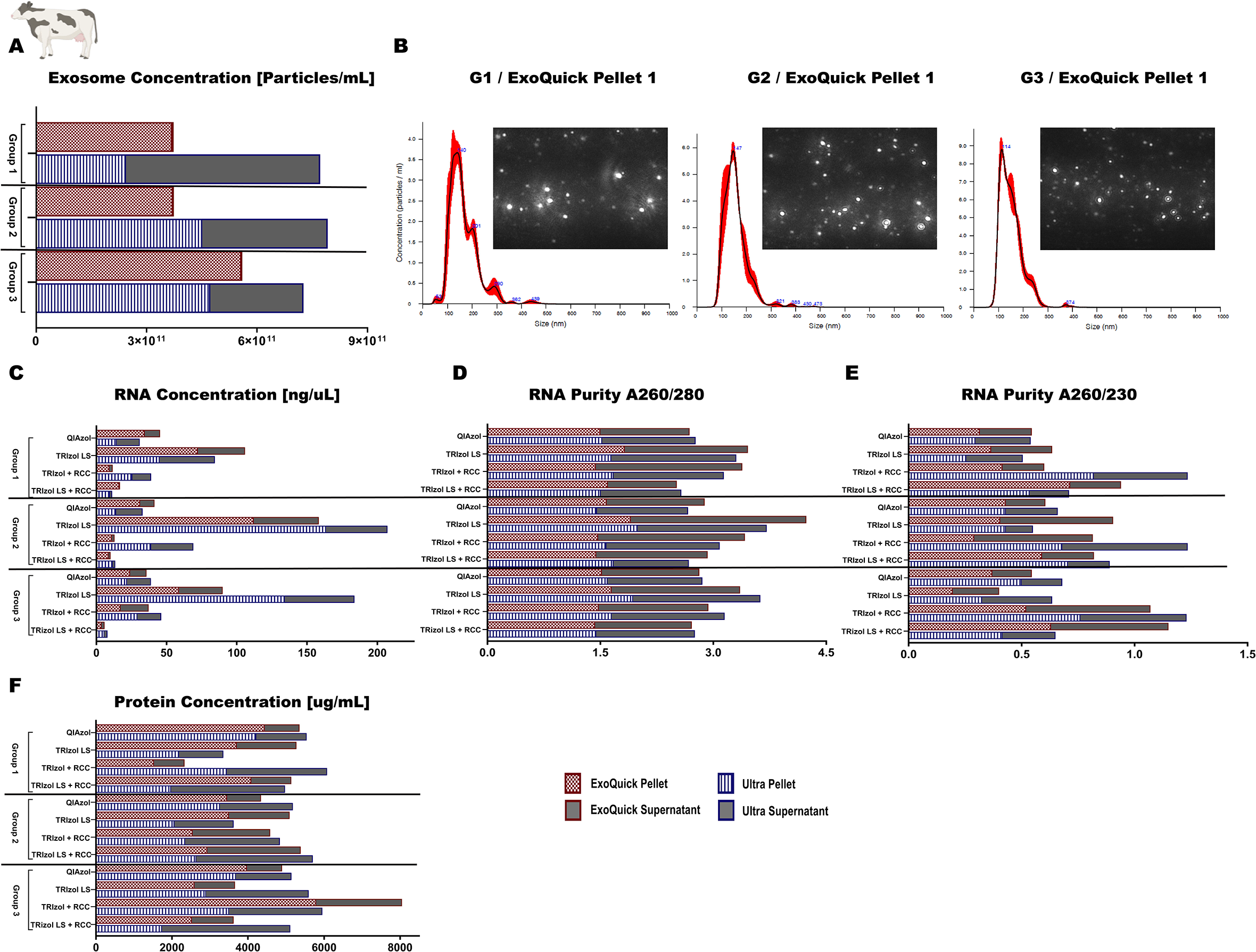
Pre-processing of whole milk prior to long-term storage of bovine milk. Group 1) unprocessed whole milk, group 2) milk without fat globules and cream, and group 3) isolated whey fraction without fat globules, cream and casein proteins. Exosome concentration [particles/mL] isolated via ExoQuick and ultracentrifugation methods across G1-G3, as determined by Nanoparticle Tracking Analysis (NTA) (A); size and distribution profiles of milk-derived exosomes as determined by NTA (B); RNA concentration [ng/μL] (C); RNA purity (A260/A280 and A260/A230) (D-E); and protein concentration [μg/mL] (F) obtained from the isolated exosomes. Data are mean ± SEM with n = 3 independent trials/group. For NTA 3 video frames of 30 s each were used. Data were analyzed using one-way analysis of variance with a Tukey post-hoc test (p ≤ 0.05) for main effect of pre-processing groups.

**Figure 2.**
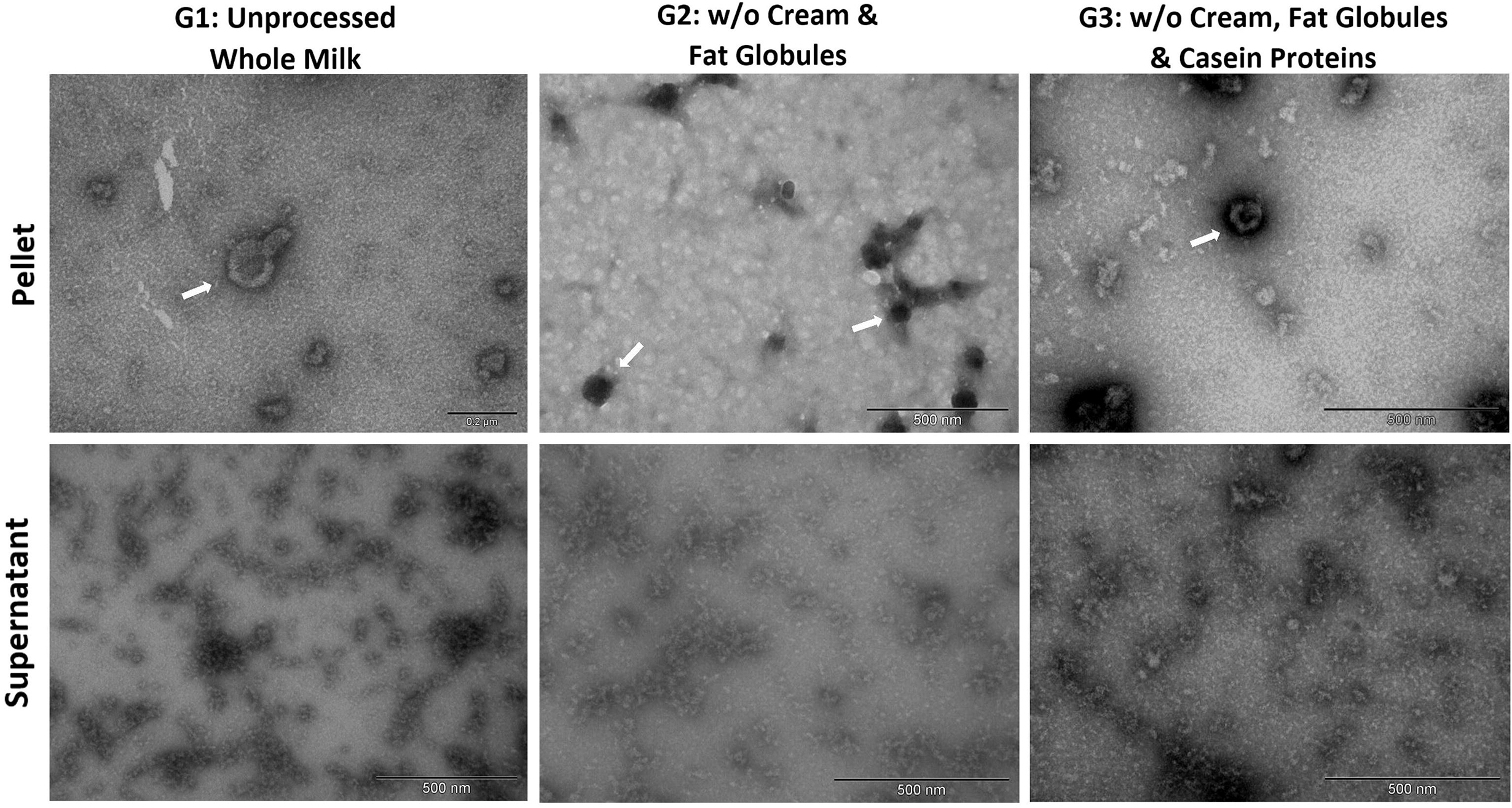
Morphology of bovine milk-derived exosomes visualized by Transmission Electron Microscopy (TEM) with negative staining (uranyl acetate). Milk-derived exosomes were isolated via ExoQuick precipitation method. Group 1) unprocessed whole milk, group 2) milk without fat globules and cream, and group 3) isolated whey fraction without fat globules, cream and casein proteins. Scale bars: 200 nm - 500 nm.

**Figure 3.**
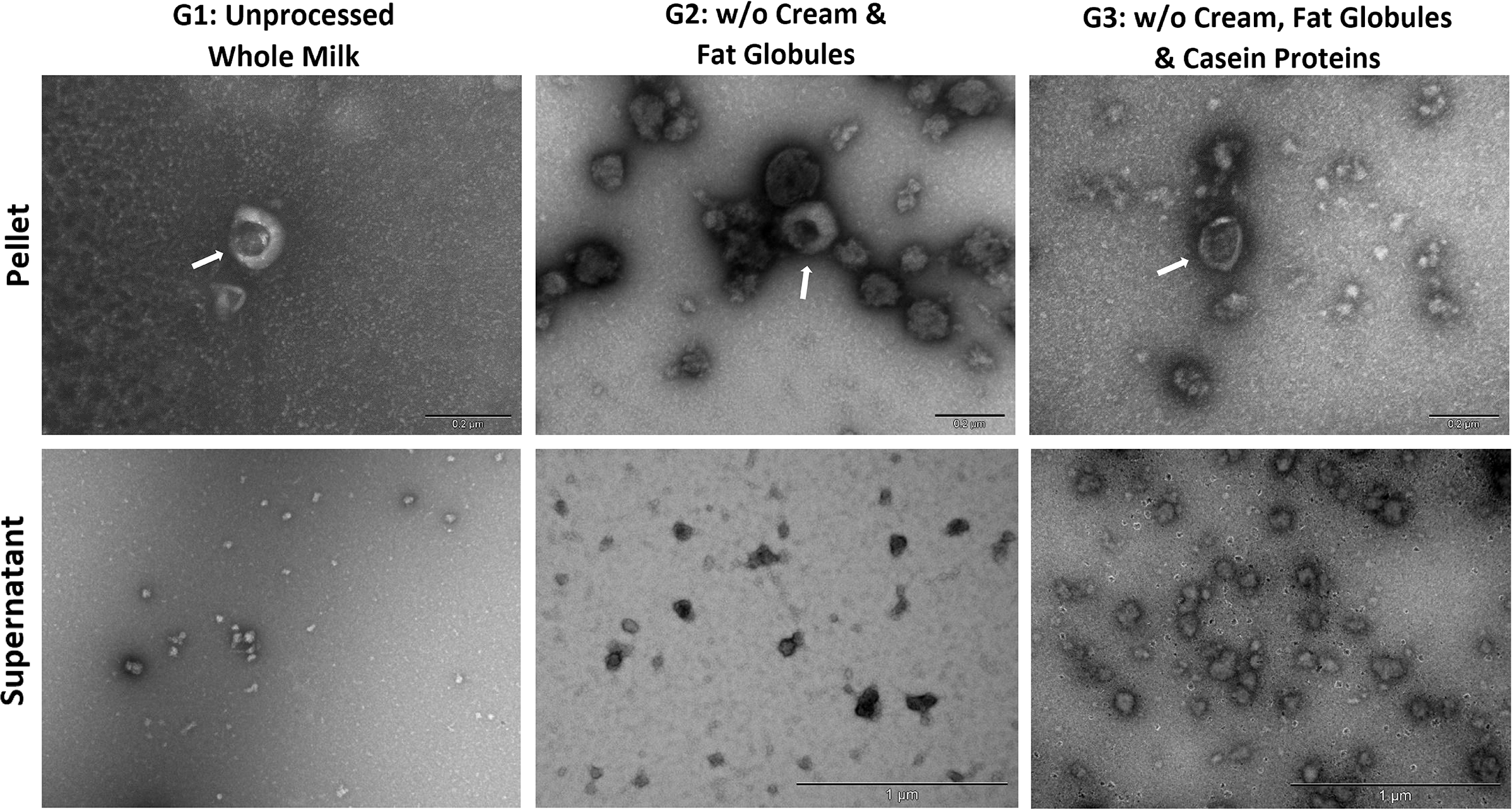
Morphology of bovine milk-derived exosomes visualized by Transmission Electron Microscopy (TEM) with negative staining (uranyl acetate). Milk-derived exosomes were isolated via ultracentrifugation method. Group 1) unprocessed whole milk, group 2) milk without fat globules and cream, and group 3) isolated whey fraction without fat globules, cream and casein proteins. Scale bars: 200 nm - 1000 nm.

### Bovine Milk-derived Exosome Isolation: EQ versus UC Methods

Bovine exosome pellets were compared against their corresponding supernatants (supernatants were used in our study as a negative control that should not contain MDEs) to test for efficiency of fractionation. According to the TEM analysis, both methods yielded more MDEs of the correct EV size (30-150nm) in the pellet fractions, when compared to their respective supernatants. Similarly, NTA indicated that exosome pellets isolated via the EQ method contained more MDEs [particles/mL], belonging to the correct particle range, when compared to the negative controls (main effect of fractionation, (*F*_(1, 23)_ = 22.236, *p* < 0.001), Tukey post hoc *p* < 0.001, Fig. 4). However, the MDE concentration between the UC pellet and the supernatant remained unchanged in bovine milk (Tukey post hoc *p* = 0.995). Additionally, EQ and UC exosome pellets resulted in higher RNA yield compared to their corresponding supernatants (main effect of fractionation, RNA concentration: (*F*_(1, 95)_ = 26.756, *p* < 0.001); A260/A280: (*F*_(1, 95)_ = 13.244, *p* < 0.001); A260/A230: (*F*_(1, 95)_ = 46.868, *p* < 0.001). Similar results were seen in gel electrophoresis, where exosome pellets extracted from both EQ and UC had visible RNA bands (ranging from 20-150 bp) compared to their respective supernatants (Fig. 5).There was also a significant difference in total soluble protein [μg/mL] between exosome pellets and their respective supernatants (*F*_(1, 95)_ = 69.884, *p* < 0.001, Fig 6)).

**Figure 4.**
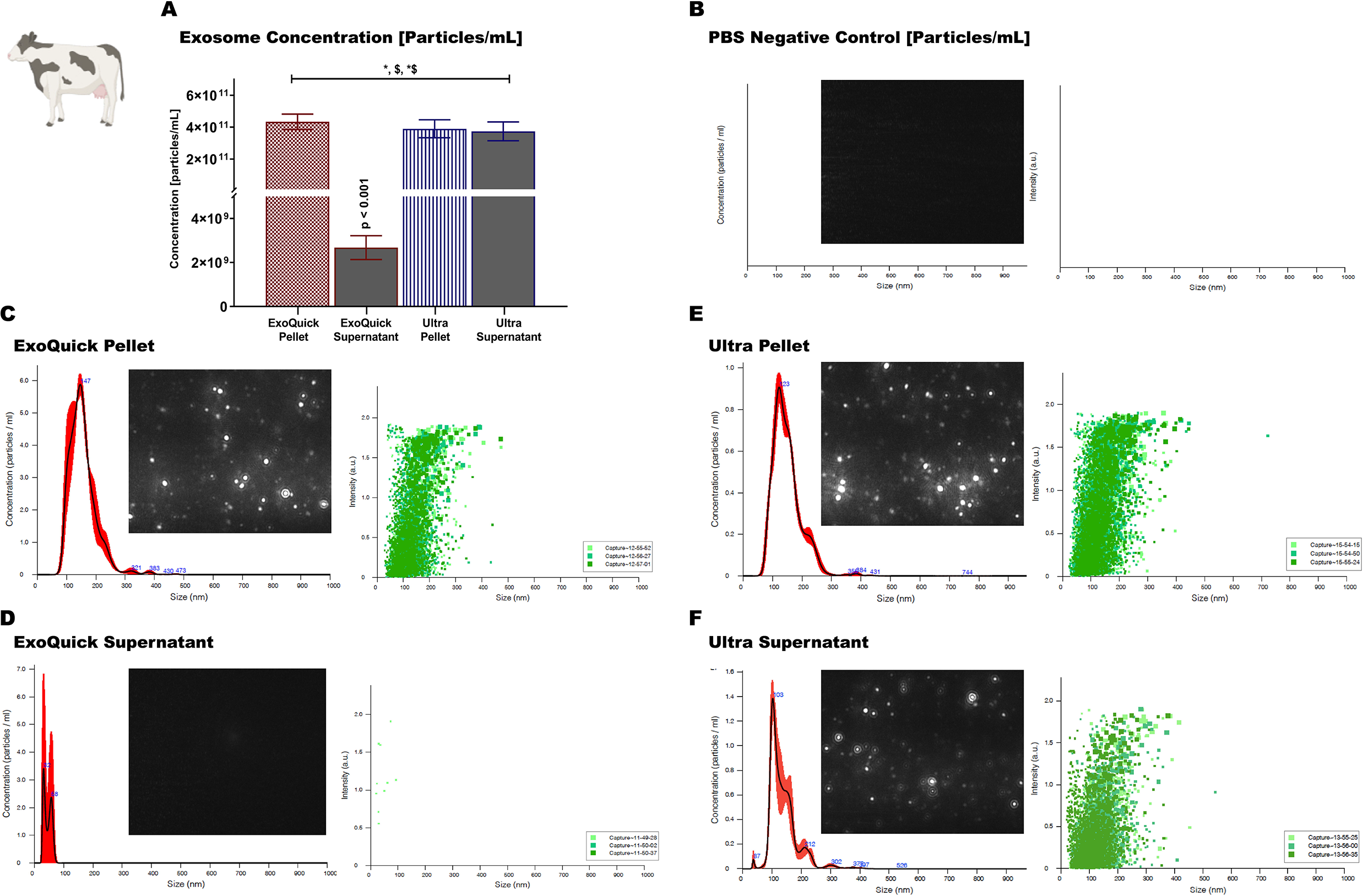
Size and distribution profiles of bovine milk-derived exosomes as determined by Nanoparticle Tracking Analysis (NTA). Exosomes were isolated via ExoQuick precipitation and ultracentrifugation methods. Exosome concentration [particles/mL] in pellets and the corresponding negative controls (the supernatants) (A). Size and distribution profiles of the absolute control (1x PBS) (B). Size and distribution profiles of the ExoQuick exosome pellet (C). Size and distribution profiles of the ExoQuick negative control (D). Size and distribution profiles of the ultracentrifugation exosome pellet (E). Size and distribution profiles of the ultracentrifugation negative control (F). Data are mean ± SEM with n = 6 independent trials/group. For NTA, 3 video frames of 30 s each were used. Data were analyzed using a two-way analysis of variance with a Tukey post-hoc test (p ≤ 0.05). Main effect of fractionation: **\***(p ≤ 0.05). Main effect of exosome isolation method: **$**(p ≤ 0.05). Fractionation/exosome isolation method interaction: **\*$** (p ≤ 0.05).

**Figure 5.**
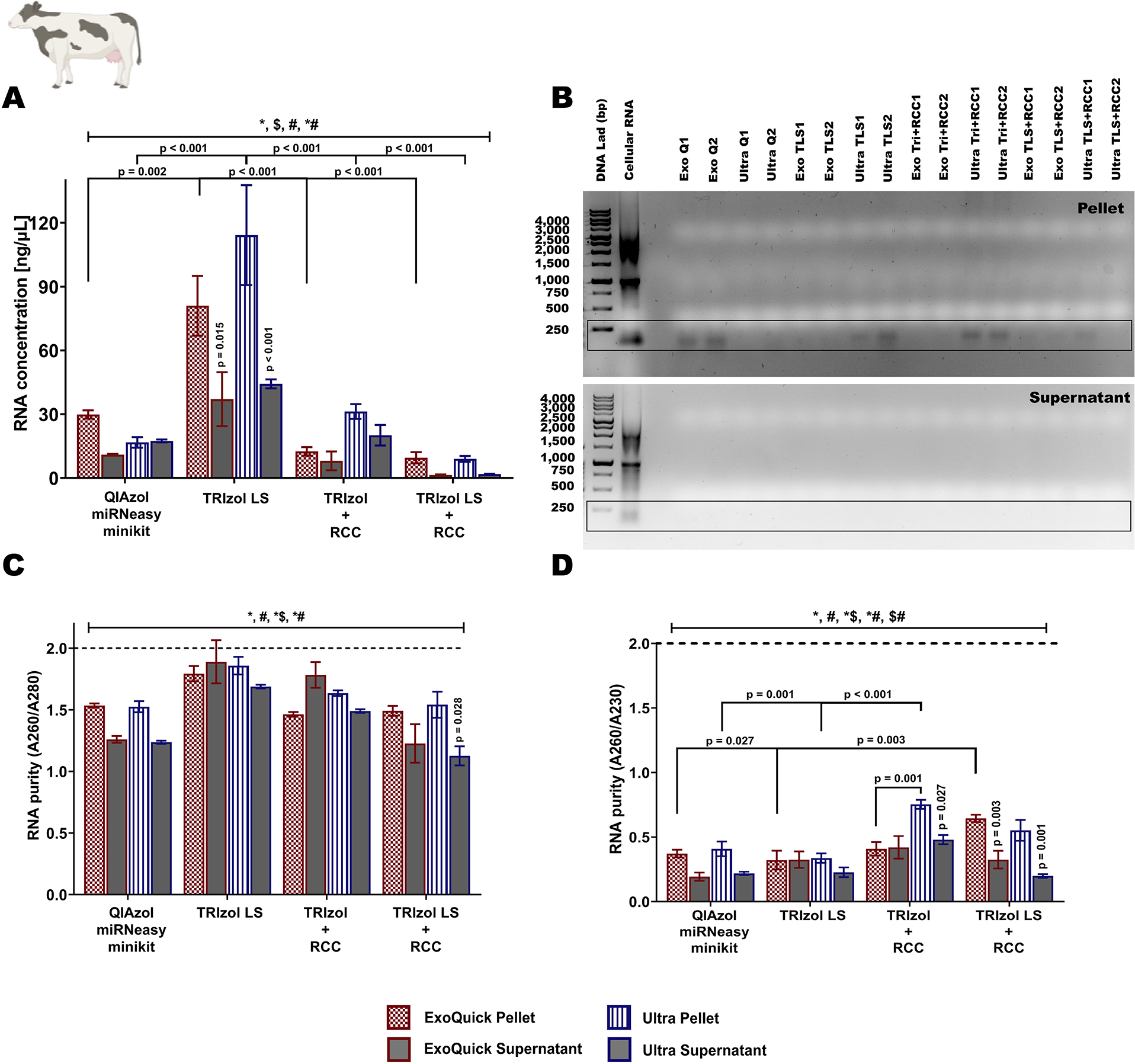
RNA yield [ng/μL], purity and quality of bovine milk-derived exosome pellets and supernatants isolated via ExoQuick precipitation and ultracentrifugation methods. RNA was extracted using four protocols, 1) QIAzol + miRNeasy MiniKit, 2) TRIzol LS, 3) TRIzol + RNA Clean and Concentrator Kit (RCC), and 4) TRIzol LS + RCC. RNA concentration [ng/μL] (A), 1 % TAE agarose gel electrophoresis (B), RNA purity - absorbance at 260nm/280nm (C), and absorbance at 260nm/230nm (D). Data are mean ± SEM with n = 6 independent trials/group. Data were analyzed using a three-way analysis of variance with a Tukey post-hoc test (p ≤ 0.05). Main effect of fractionation: **\***(p ≤ 0.05). Main effect of exosome isolation method: **$**(p ≤ 0.05). Main effect of RNA extraction protocol: **#**(p ≤ 0.05). Fractionation/exosome isolation interaction: **\*$** (p ≤ 0.05). Fractionation/RNA extraction interaction: **\*#**(p ≤ 0.05). Exosome isolation/RNA extraction interaction: **$#**(p ≤ 0.05). p-values on top of supernatant bars indicate significant difference between pellets and the corresponding supernatants of that particular exosome isolation protocol and RNA extraction protocol.

**Figure 6.**
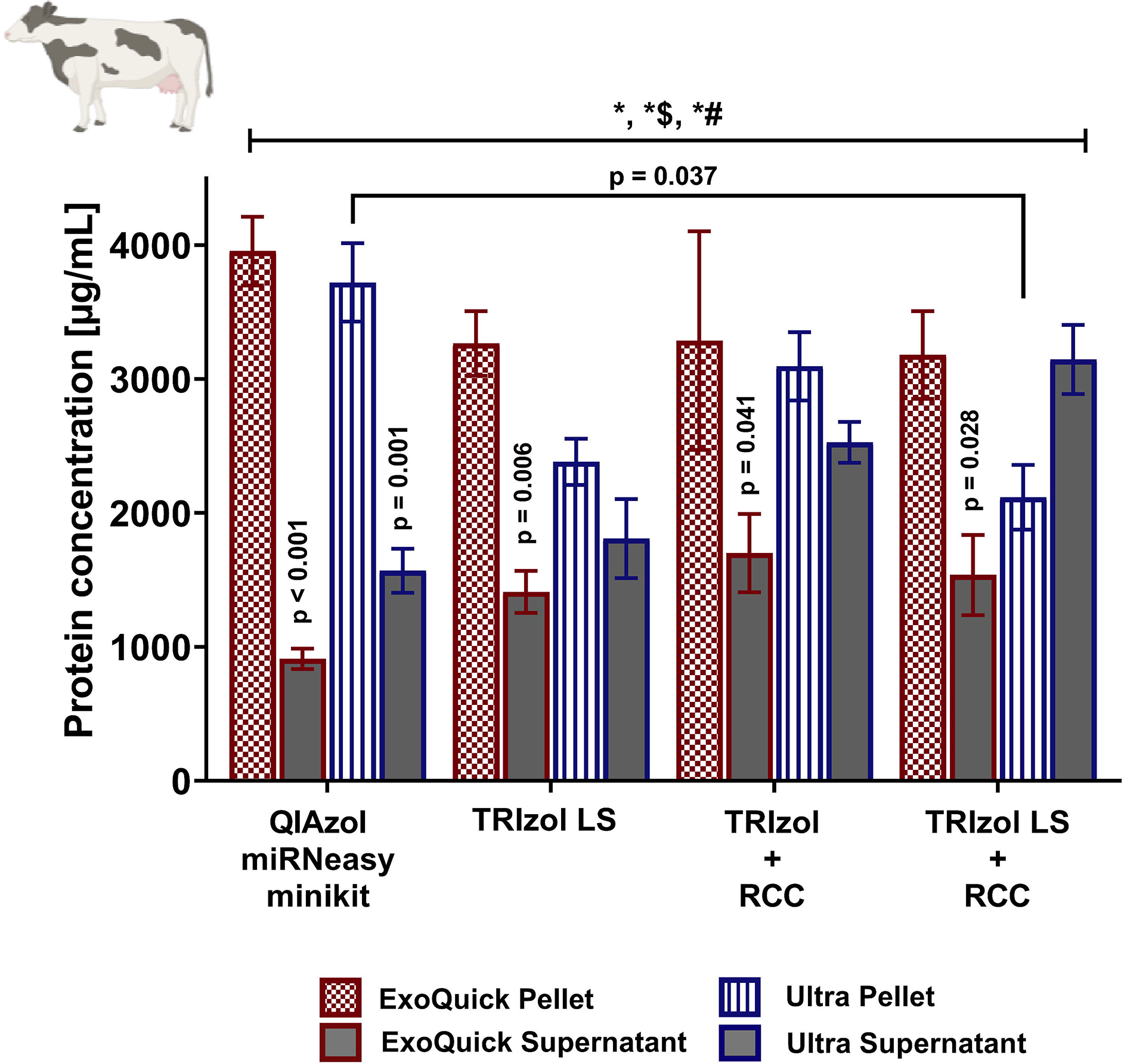
Protein concentration [μg/mL] of bovine milk-derived exosome pellets and supernatants isolated via ExoQuick precipitation and ultracentrifugation methods. Total soluble proteins were extracted from the lower organic phase resulting from four RNA extraction protocols, 1) QIAzol + miRNeasy MiniKit, 2) TRIzol LS, 3) TRIzol + RNA Clean and Concentrator Kit (RCC), and 4) TRIzol LS + RCC. Data are mean ± SEM with n = 6 independent trial/group. Data were analyzed using a three-way analysis of variance with a Tukey post-hoc test (p ≤ 0.05). Main effect of fractionation: **\***(p ≤ 0.05). Fractionation/exosome isolation interaction: **\*$** (p ≤ 0.05). Fractionation/RNA extraction interaction: **\*#**(p ≤ 0.05). p-values on top of supernatant bars indicate significant difference between pellets and the corresponding supernatants of that particular exosome isolation protocol and RNA extraction protocol.

EQ versus UC isolation methods were compared to determine whether one method is more suitable than the other for isolating high quality, intact MDEs. According to TEM results, exosome pellets isolated via UC contained more intact, spherical exosomes of the correct particle size (30-150nm) compared to the EQ pellets. However, the NTA results indicated that EQ pellets contained a higher concentration of exosomes of the correct particle size compared to UC (main effect of exosome isolation method, (*F*_(1, 23)_ = 11.942, *p* < 0.002)). Minimal differences were seen in RNA concentration, RNA purity (A260/A80; A260/A230), and protein concentration [μg/mL] of exosome pellets obtained via EQ and UC methods (main effect of exosome isolation method, A260/A280: (*F*_(1, 95)_ = 1.186, *p* = 0.279); A260/A230: (*F*_(1, 95)_ = 0.629, *p* = 0.430); and protein concentration: (*F*_(1, 95)_ = 0.814, *p* = 0.370)).

### RNA Extraction of Bovine Milk-derived Exosomes

The TRIzol LS protocol produced the highest RNA yield [ng/μL] for bovine MDEs isolated via EQ and UC methods. EQ: TRIzol LS vs QIAzol miRNeasy minikit (main effect of RNA extraction protocol, (*F*_(3, 95)_ = 51.215, *p* < 0.001), Tukey post hoc *p* = 0.002), TRIzol LS vs TRIzol + RCC (Tukey post hoc *p* < 0.001), and TRIzol LS vs TRIzol LS + RCC (Tukey post hoc *p* < 0.001). UC: TRIzol LS vs QIAzol miRNeasy minikit (Tukey post hoc *p* < 0.001), TRIzol LS vs TRIzol + RCC (Tukey post hoc *p* < 0.001), and TRIzol LS vs TRIzol LS + RCC (Tukey post hoc *p* < 0.001). There was also a significant fractionation/RNA extraction interaction (*F*_(3, 95)_ = 9.569, *p* < 0.001).

RNA purity (A260/A280) showed a significant main effect of RNA extraction protocol, (*F*_(3, 95)_ = 29.122, *p* < 0.001) and fractionation/RNA extraction interaction, (*F*_(3, 95)_ = 6.637, *p* < 0.001). Moreover, TRIzol + RCC and TRIzol LS + RCC protocols produced higher quality RNA (A260/A230) compared to QIAzol miRNeasy minikit and TRIzol LS protocols. UC exosome pellets: TRIzol + RCC vs QIAzol miRNeasy minikit (main effect of RNA extraction protocol, (*F*_(3, 95)_ = 16.805, *p* < 0.001), Tukey post hoc *p* = 0.001), TRIzol + RCC vs TRIzol LS (Tukey post hoc *p* < 0.001). EQ exosome pellets: TRIzol LS + RCC vs QIAzol miRNeasy minikit (Tukey post hoc *p* = 0.027), and TRIzol LS + RCC vs TRIzol LS (Tukey post hoc *p* = 0.003).

Total soluble protein [μg/mL] isolated from the lower organic-phase of the four phenol-based RNA extractions remained unchanged across methods (main effect of RNA extraction protocol (*F*_(3, 95)_ = 1.416, *p* = 0.244)).

### Human Milk-derived Exosome Isolation: EQ versus UC Methods

Human MDEs isolated via EQ and UC methods were compared against their corresponding supernatants to determine the efficiency of fractionation. TEM results confirmed that EQ and UC pellets contained more intact exosomes of the correct MV size compared to their respective supernatants (Fig. 7). NTA analysis also indicated that exosome pellets isolated via both EQ and UC methods produced more exosomes [particles/mL] compared to their corresponding supernatants (main effect of fractionation, (*F*_(1, 7)_ = 534.670, *p* < 0.001), Tukey post hoc *p* < 0.001, Fig. 8). Similarly, RNA concentration [ng/μL] (*F*_(1, 31)_ = 131.638, *p* < 0.001) and RNA purity (A260/A280 (*F*_(1, 31)_ = 154.293, *p* < 0.001); A260/A230 (*F*_(1, 31)_ = 111.250, *p* < 0.001)) of the exosome pellets were higher than their respective supernatants. Corresponding results were seen in the 1% TAE agarose gel, where exosomes pellets isolated via EQ and UC methods had visible RNA bands (ranging from 20-200 base pairs) compared to their supernatants (Fig. 9). Total soluble protein concentration [μg/mL] was also higher in the pellet fractions (*F*_(1,31)_ = 28.778, *p* < 0.001, Fig. 10) compared to the supernatants.

**Figure 7.**
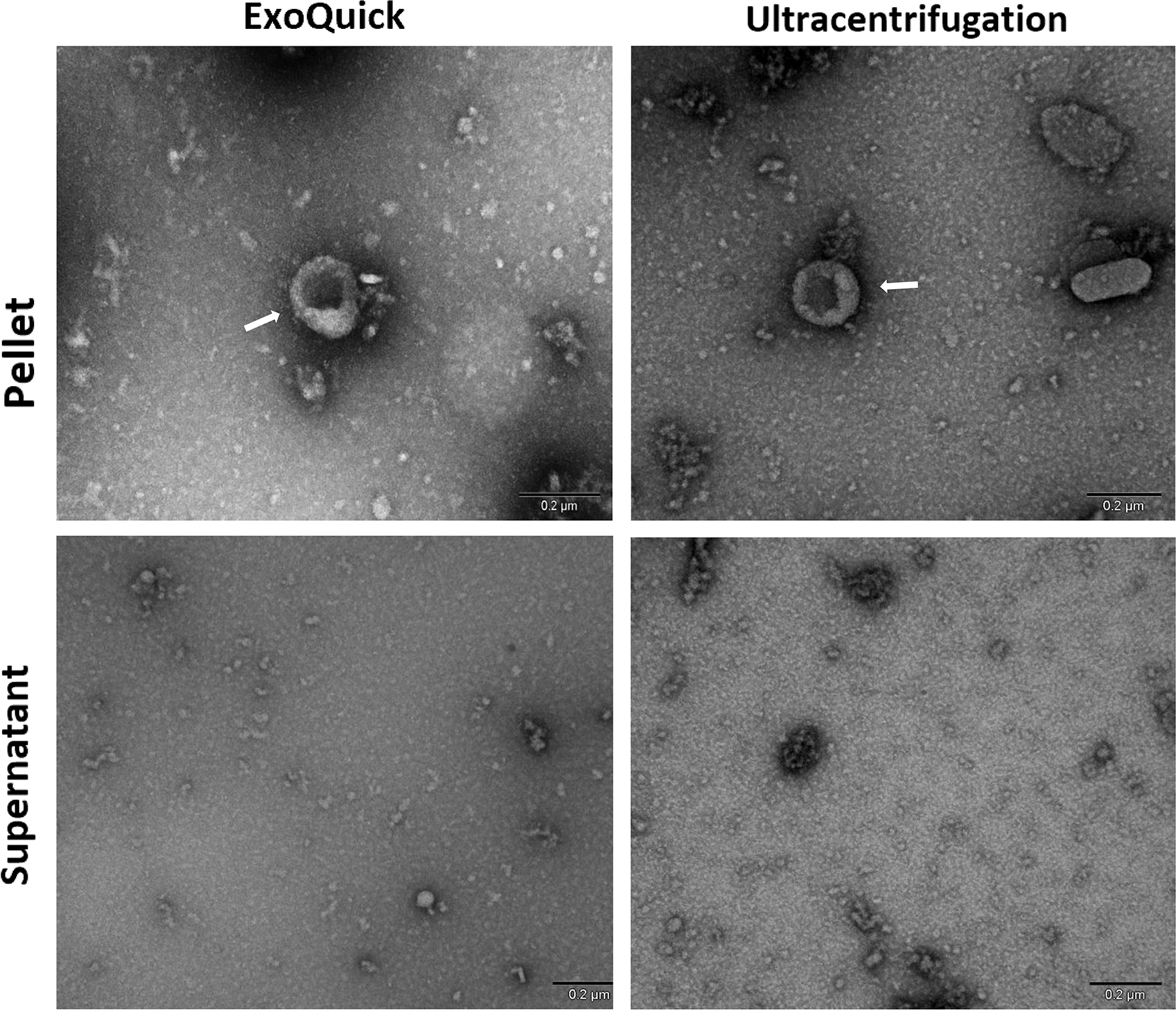
Morphology of human milk-derived exosomes visualized by Transmission Electron Microscopy (TEM) with negative staining (uranyl acetate). Human milk-derived exosomes were isolated via ExoQuick precipitation and ultracentrifugation methods. Scale bars: 200 nm.

**Figure 8.**
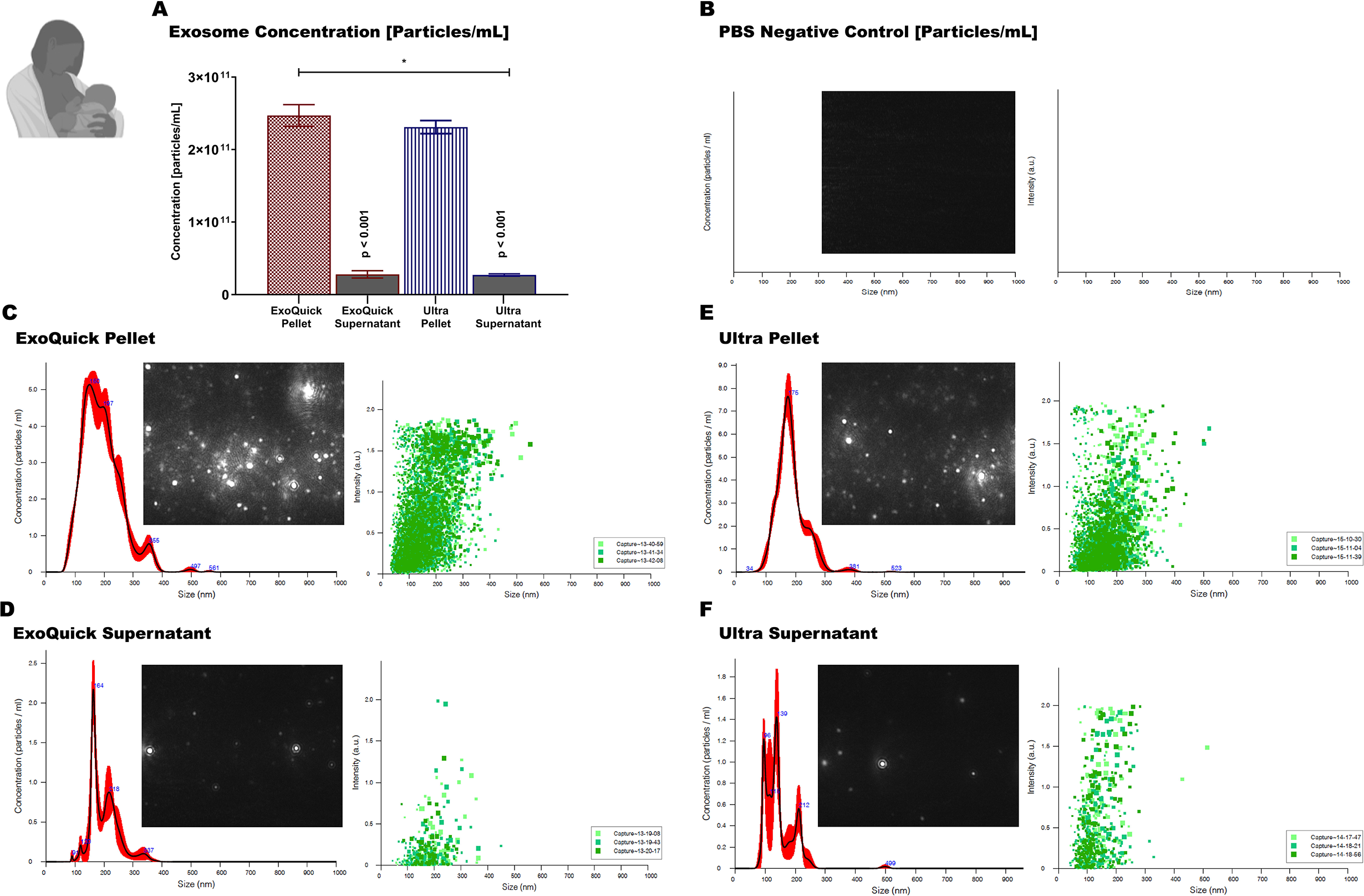
Size and distribution profiles of human milk-derived exosomes as determined by Nanoparticle Tracking Analysis (NTA). Exosomes were isolated via ExoQuick precipitation and ultracentrifugation methods. Exosome concentration [particles/mL] of the pellet and the corresponding negative controls (the supernatants) (A). Size and distribution profiles of the absolute control (1x PBS) (B). Size and distribution profiles of the ExoQuick exosome pellet (C). Size and distribution profiles of the ExoQuick negative control (D). Size and distribution profiles of the ultracentrifugation exosome pellet (E). Size and distribution profiles of the ultracentrifugation negative control (F). Data are mean ± SEM with n = 6 independent trials/group. For NTA, 3 video frames of 30 s each were used. Data were analyzed using a two-way analysis of variance with a Tukey post-hoc test (p ≤ 0.05). Main effect of fractionation: **\***(p ≤ 0.05). Main effect of exosome isolation method: **$**(p ≤ 0.05). Fractionation/exosome isolation method interaction: **\*$** (p ≤ 0.05).

**Figure 9.**
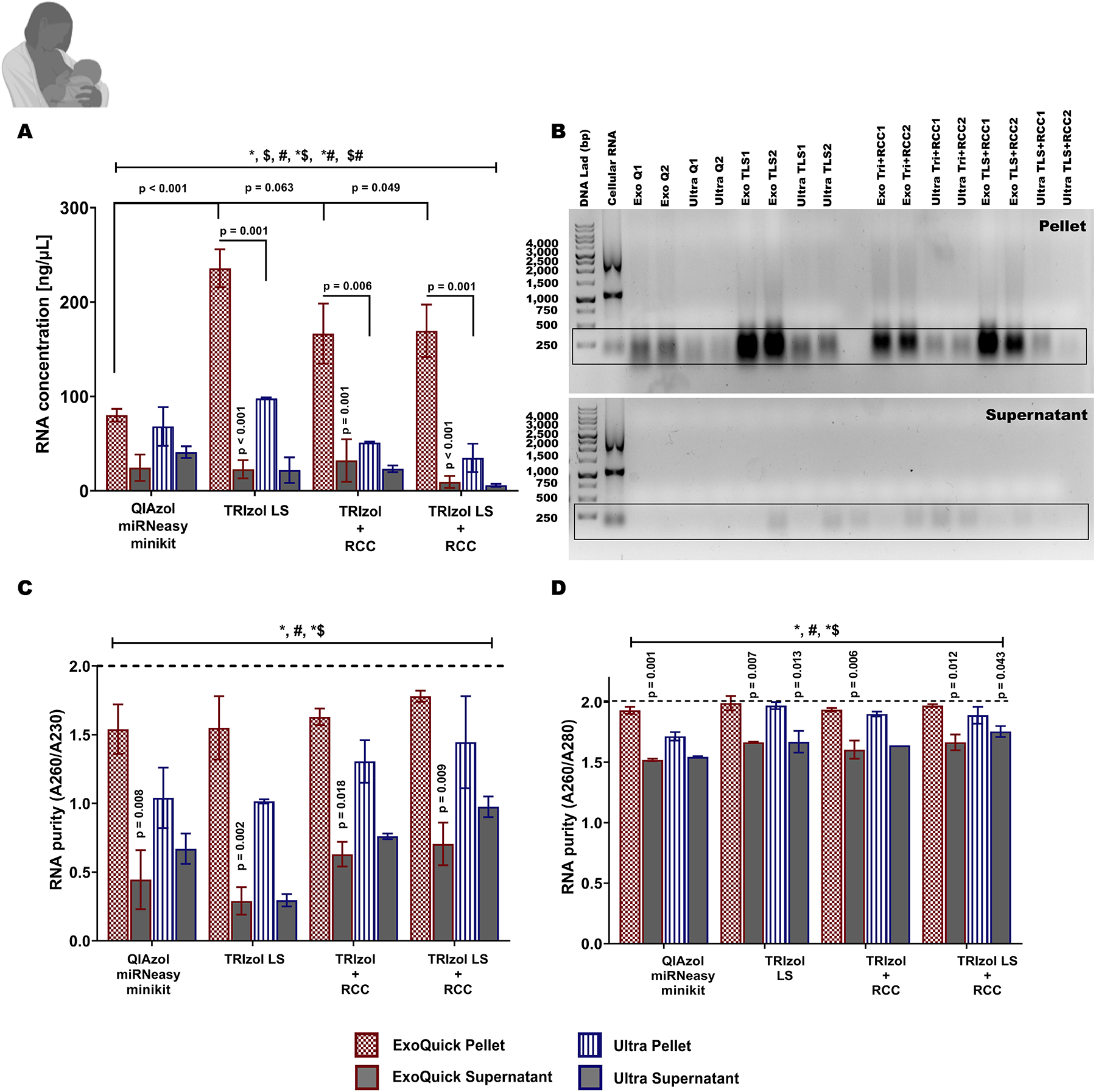
RNA yield [ng/μL], purity and quality of human milk-derived exosome pellets and supernatants isolated via ExoQuick precipitation and ultracentrifugation methods. RNA was extracted using four protocols, 1) QIAzol + miRNeasy MiniKit, 2) TRIzol LS, 3) TRIzol + RNA Clean and Concentrator Kit (RCC), and 4) TRIzol LS + RCC. RNA concentration [ng/μL] (A), 1 % TAE agarose gel electrophoresis of exosome RNA (B), RNA purity - absorbance at 260nm/280nm (C), and absorbance at 260nm/230nm (D). Data are mean ± SEM with n = 3 technical replicates/exosome isolation method /RNA extraction protocol. Data were analyzed using a three-way analysis of variance with a Tukey post-hoc test (p ≤ 0.05). Main effect of fractionation: **\***(p ≤ 0.05). Main effect of exosome isolation method: **$**(p ≤ 0.05). Main effect of RNA extraction protocol: **#**(p ≤ 0.05). Fractionation/exosome isolation interaction: **\*$** (p ≤ 0.05). Fractionation/RNA extraction interaction: **\*#**(p ≤ 0.05). Exosome isolation/RNA extraction interaction: **$#**(p ≤ 0.05). p-values on top of supernatant bars indicate significant difference between pellets and the corresponding supernatants of that particular exosome isolation protocol and RNA extraction protocol.

**Figure 10.**
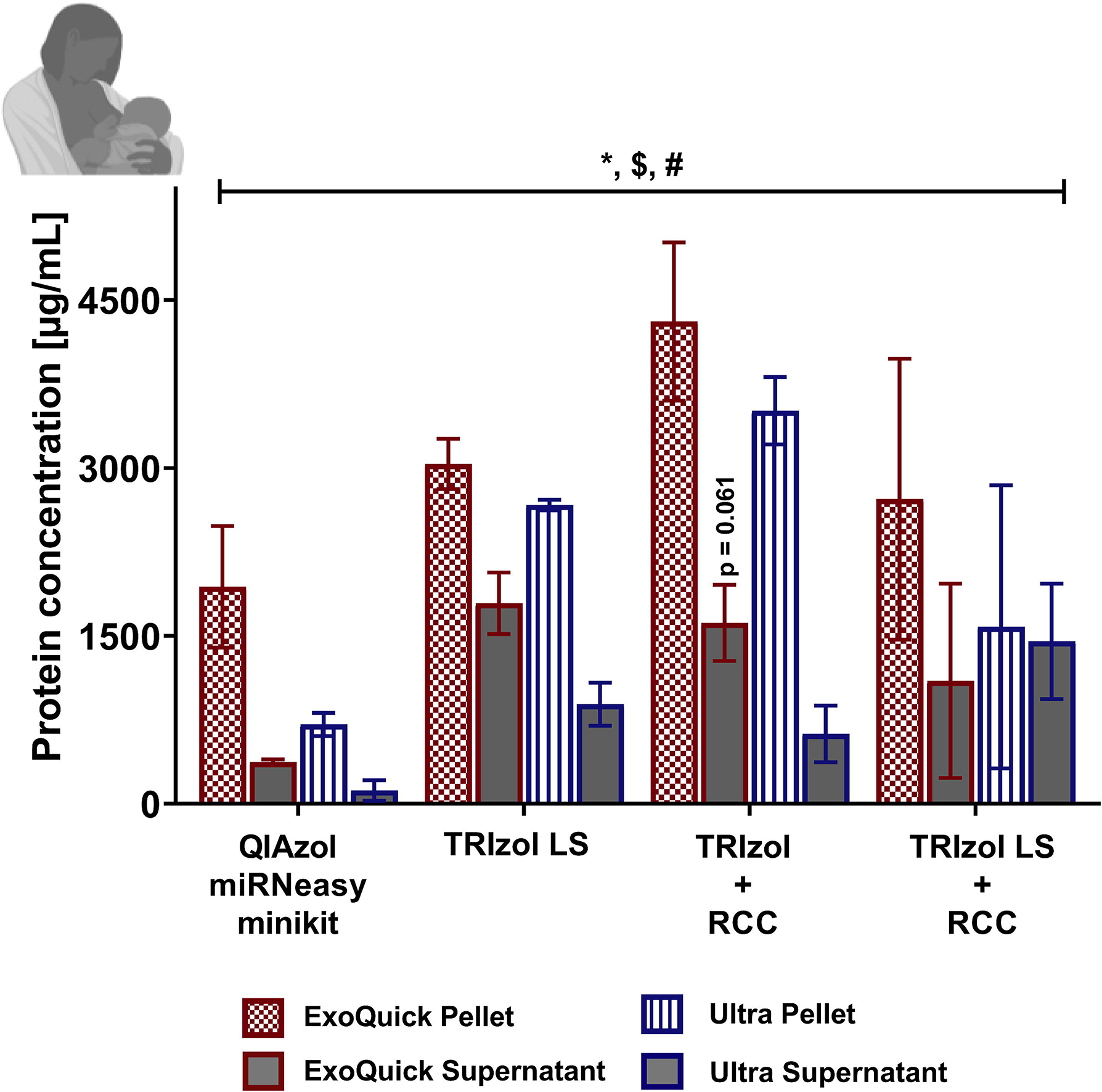
Protein concentration [μg/mL] of human milk-derived exosome pellets and supernatants isolated via ExoQuick precipitation and ultracentrifugation methods. Total soluble proteins were extracted from the lower organic phase resulting from four RNA extraction protocols, 1) QIAzol + miRNeasy MiniKit, 2) TRIzol LS, 3) TRIzol + RNA Clean and Concentrator Kit (RCC), and 4) TRIzol LS + RCC. Data are mean ± SEM with n = 6 independent trial/group Data were analyzed using a three-way analysis of variance with a Tukey post-hoc test (p ≤ 0.05). Main effect of fractionation: **\***(p ≤ 0.05). Main effect of exosome isolation method: **$**(p ≤ 0.05). Main effect of RNA extraction protocol: **#**(p ≤ 0.05). p-values on top of supernatant bars indicate significant difference between pellets and the corresponding supernatants of that particular exosome isolation protocol and RNA extraction protocol.

According to the TEM, the EQ pellets contained less cell debris and more intact exosomes within the expected size range (30-150 nm) compared to the UC exosome pellets. However, the NTA analysis indicated no differences in exosome concentration [particles/mL] between EQ and UC isolation methods (main effect of exosome isolation method, (*F*_(1, 7)_ = 0.848, *p* = 0.409)). EQ exosome pellets contained more RNA [ng/μL] (main effect of exosome isolation method, (*F*_(1, 31)_ = 39.609, *p* < 0.001)), and total soluble protein [μg/mL] (*F*_(1, 31)_ = 5.220, *p* = 0.036) than UC exosome pellets. There was a significant fractionation/exosome isolation interaction (RNA concentration: (*F*_(1, 31)_ = 40.938, *p* < 0.001); A260/A280: (*F*_(1, 31)_ = 7.877, *p* = 0.013); and A260/A230: (*F*_(1, 31)_ = 14.082, *p* = 0.002)).

### RNA Extraction of Human Milk-derived Exosome

Minimal differences were seen across TRIzol LS, TRIzol + RCC, and TRIzol LS + RCC extractions methods, however the QIAzol miRNeasy minikit produced the lowest RNA yield [ng/μL] compared to the other protocols; TRIzol LS (main effect of RNA extraction protocol, (*F*_(3, 31)_ = 5.862, *p* = 0.007), Tukey post hoc *p* < 0.001), TRIzol + RCC (Tukey post hoc *p* = 0.063), and TRIzol LS + RCC (Tukey post hoc *p* = 0.049). There was a significant fractionation/RNA extraction interaction (*F*_(3, 31)_ = 7.286, *p* = 0.003) and exosome isolation/RNA extraction interaction (*F*_(3, 31)_ = 4.855, *p* = 0.014). There was also significant main effect of RNA extraction protocol on RNA purity (A260/A280: (*F*_(3, 31)_ = 9.168, *p* = 0.001); A260/A230: (*F*_(3, 31)_ = 6.039, *p* = 0.006)) and protein concentration: (*F*_(3, 31)_ = 6.413, *p* = 0.005).

## Discussion

Recent studies have identified a number of novel bioactive components in maternal milk, including lactation-specific miRNAs and stem cells. The abundance of lactation-specific miRNAs in milk fractions (17,18,21,50), their stability in offspring’s GI tracts (11,13,16,41), and their localization in peripheral organs (14), the blood (19), and the brain (24), suggest an important role for maternal milk miRNAs in postnatal development. Reproducible and standardized methods for milk processing, storage, and MDE isolation are required to ensure accurate characterization of MDEs and their function. In this study, we tested three milk pre-processing methods, two commonly used MDE isolation techniques, and four RNA extraction protocols used to obtain high quality exosomal RNA from bovine and human milk. Collectively, our results indicate that pre-processing of whole milk to remove fat, cream, and casein proteins prior to long-term storage is not required to obtain high-quality MDEs. Moreover, EQ precipitation is better suited for the isolation of exosomes from bovine and human whey milk fractions compared to UC. Further, the TRIzol LS protocol produced the highest RNA yield from bovine MDEs, while TRIzol LS, TRIzol+RCC, and TRIzol LS+RCC methods were found to be efficient for the extraction of high quality human MDEs.

### Pre-processing of Bovine Milk

Some studies have reported that the pre-processing of whole milk to remove contaminants and unwanted components prior to long-term storage affects the yield and the purity of isolated MDEs (29,31), while other studies have reported minimal effects of pre-processing on the quality of MDEs (12,51,52). We found some differences in exosome quality among the three pre-processing groups via TEM. However, NTA, RNA, and protein data confirmed that the pre-processing of milk prior to long-term storage at −80 °C is not necessary to obtain high quality MDEs (Fig. 1). The visual differences in TEM images that were recorded across G1-G3 are not surprising given that operator image selection can lead to false interpretations regarding EV quality. As others have shown, operator image selection is more suitable to demonstrate the presence of EVs in a sample, but is less suitable to demonstrate the quality of the fractionation (29). Our results are thus consistent with other published studies indicating that pre-processing of whole milk prior to long-term storage does not affect the final MDE yield, integrity and/or biological activities (51,52). Further, hold pasteurization involving 62.5 °C temperature treatment for 30 min also does not appear to adversely affect the integrity and biological function of MDEs, and pasteurized MDEs were shown to be as functionally beneficial as raw milk MDEs in therapeutics (12). However, a few studies have shown the opposite results, where pre-processing of whole milk prior to storage was required for successful MDE isolations (29,30,51). Notably, these select studies used different storage conditions, handling procedures, sample preparation, and exosome isolation techniques compared to the methods used in our study. In particular, Zonneveld et al. (2014) processed a batch of human milk immediately after collection, processed two more batches after storing at −80 °C and 4 °C for 2 h, and used density gradient centrifugation for isolating MDEs (29). In contrast, the samples of human donor milk used in our study were obtained from a human milk bank and it was stored immediately upon collection at - 20 °C and thawed at 4 °C overnight before processing. Note: the standard procedure for the storage of human milk by milk banks is that all human milk samples are frozen immediately upon collection by the donors at −20 °C, transported to the milk bank in a frozen state, and subsequently stored at −20 °C until ready for pasteurization and processing. As such, differences in sample handling conditions limit direct comparisons between these studies and the present investigation. This highlights one of the main limitations of current MDE research, where a lack of standardized methods for sample handling/storage and MDE isolations may lead to erroneous conclusions and incomparable results. Our findings suggest that it is feasible to process whole milk after long-term storage without incurring significant losses in MDE concentration, purity, and integrity. This is a very useful finding as all donor milk obtained from human milk banks is frozen soon after expression without pre-processing. As a majority of MDE research in human milk involves samples obtained from milk banks, the ability to isolate high quality MDEs in large quantities post-long-term storage is of great value. In addition, there is evidence that eliminating time sensitive pre-processing requirements may also increase the viability of other bioactive and macro/micronutrient components of milk, including hormones, vitamins, and growth factors that are sensitive to repeated handling and temperature fluctuations (29,31).

### Bovine Milk-derived Exosome Isolation: UC versus EQ Methods

UC-based isolations, EQ precipitation, size-based isolations, and immunity capture-based techniques are some of the most commonly used exosome isolation methods. Each technique utilizes a particular trait of EVs, including density, shape, size, and/or surface receptors/membrane proteins for the isolations. As such, there are unique advantages and disadvantages to each method (33). A robust method with the ability to minimize exosome loss, lysis, as well as co-isolation of contaminants is an important pre-requisite that should be standardized across MDE research in order to minimize confounding effects that can alter downstream applications. Density gradient centrifugation, where exosomes are separated based on size, mass, and density in a sucrose gradient, is the gold standard for exosome isolations. However, an important drawback of this technique is the loading limitation, where only a small volume of milk must be loaded onto the gradient (33). Density gradient centrifugation can also be labor-intensive and can take more than 24 h to complete (53). Thus, differential UC (100,000 x g for 90 min) coupled with serial filtration has become increasingly popular for large-scale MDE isolations in recent years (12,37,38,44). However, due to the mechanical stress of ultra-high speeds and increased handling that is associated with each step, UC can suffer from exosome loss (54). We also found that the UC method resulted in a large loss of MDEs in bovine milk (Fig. 4). In particular, the UC-supernatant (the designated negative control) contained a similar concentration of MDEs compared to the enriched pellet, indicative of an unsuccessful fractionation and a large loss of MDEs. Specific g-force /k factor usage and centrifugation run times during UC may have contributed to this, as speeds of 100,000 x g or more can greatly influence the sedimentation ratio of pure exosomes (55). Serial UC methods may need to be coupled with another purification protocol, including density gradient centrifugation and/or buoyant density gradient centrifugation to isolate high quality MDEs from bovine milk whey (12,26,33,37,50,56). Nevertheless, the EQ method successfully fractionated MDEs in bovine milk, where a significant difference in exosome concentration was observed between the pellet and the supernatant. We found that EQ precipitation is more effective at isolating MDEs from bovine milk whey compared to the UC method. However, minimal differences were seen in the total RNA (Fig. 5) and soluble protein (Fig. 6) obtained between the two methods. It is therefore imperative to consider the downstream molecular applications that are associated with the MDE isolations prior to choosing a method. Notably, this is also one of the major recommendations of ISEV2014 (28).

### Human Milk-derived Exosome Isolation: UC versus EQ Methods

In human milk, both the UC and EQ methods successfully isolated MDEs, as evidenced by the higher concentrations of exosomes (Fig. 7–8), RNA (Fig. 9), and protein (Fig. 10) in the pellet fractions compared to their respective supernatants. However, similar to bovine milk, we found the EQ method to be more efficient at isolating high quality intact MDEs of the correct size from human milk compared to the UC method. Indeed, EQ has been shown to be far superior than size exclusion, immunoaffinity, and UC methods by previous studies, as the polymer-based technology alters the solubility and dispersibility of the biological fluid and force insoluble EVs out of the solution (32,33,42,57). Similarly, several maternal milk-based studies to date have used EQ method to successfully isolate MDEs across bovine, human, and rodent milk (13,38,56,58). It should be noted that EQ precipitation requires pre-clean up steps and may also co-precipitate non-exosome contaminants, including protein aggregates and other unwanted MVs (33,54). As such, in order to accurately compare UC and EQ precipitation methods, in terms of exosome yield, purity, and size exclusivity, we added several thorough pre-purification steps to remove cellular debris, fat, cream, and casein proteins, and serial filtration steps (0.45 μM and 0.22 μM) to both methods.

### RNA Extraction of Bovine and Human Milk-derived Exosomes

High RNA yield, purity, and integrity are essential parameters that can determine the validity of MDE-based molecular applications, including RT-qPCR, microarrays, RNA/miRNA sequencing. We found that the TRIzol LS protocol is best suited for extracting RNA from bovine MDEs. Similarly, previous studies have also isolated high quality RNA, suitable for RNA sequencing, from bovine MDEs using the Trizol LS protocol (47,59). Moreover, we did not detect significant differences in human MDE RNA yield and purity across TRIzol LS, TRIzol + RCC, and TRIzol LS + RCC protocols. We concluded that all three methods can be used to isolate sequencing-quality exosome RNA from human MDEs. The QIAzol + miRNeasy Mini Kit is the least preferable RNA extraction method, as this protocol produced the lowest RNA yield. Although several previous studies have reported successful RNA extractions using QIAzol + miRNeasy Mini Kit in bovine, mice, and human milk samples, their sample processing, exosome extraction techniques, starting material, and parturient histories varied from ours (11,15,21,45,48). For example, Oh et al. (2015) stored bovine colostrum immediately after collection at −80 °C until further use and only conducted differential centrifugations to remove fat and casein proteins. Whereas, Izumi et al. (2012) stored bovine colostrum immediately at −20 °C after collection and conducted serial filtrations to eliminate residual cell debris along with removing fat and casein proteins via differential centrifugation. Our samples were not stored at - 80 °C and/or −20 °C immediately upon collection, and we performed serial filtration and processing steps prior to MDE isolations via the UC method.

In conclusion, our findings indicate that it is not necessary to pre-process unpasteurized bovine milk to remove cream, fat globules, and casein proteins soon after collection and prior to long-term storage at −80 °C. Similar quantities and quality of MDEs can be obtained from frozen whole milk, as long as the processing to remove unwanted materials is conducted prior to MDE fraction. Furthermore, we have refined an EQ method, coupled with serial centrifugation and filtration steps, that can be used to successfully precipitate high quality MDEs from both bovine and human frozen whole milk. We also identified TRIzol LS as best suited for the isolation of bovine MDE RNA, whereas TRIzol LS, TRIzol + RCC, and TRIzol LS + RCC methods can be used interchangeably in human samples. Taken together, our findings provide new experimental insights into MDE processing and storage requirements, fractionation techniques, RNA, and protein extraction methods that can be used to establish standardized guidelines and protocols for MDE isolation, characterization, and analysis. Standardized protocols will enhance the comparability and reproducibility of MDE research and minimize confounding variables that may mask relevant biological differences across milk samples.

## Acknowledgements

We like to thank Loa-De-Mede Holsteins Farm in Oshawa, Ontario, Canada for providing unpasteurized, fresh, bovine milk. We like to thank Ms. Durga Archarya and Mr. Bruno Chue at the Center for the Neurobiology of Stress, Toronto, Ontario, Canada for their valuable assistance and guidance with Transmission Electron Microscopy. We like to thank Mr. Greg Wesney and Mr. James Jorgensen at The Structural & Biophysical Core Facility at the Peter Gilgan Center for Research and Learning, Toronto, Ontario, Canada for assistance with Nanoparticle Tracking Analysis. We like to also thank Dr. Deborah O’Connor and Dr. Agostino Pierro at the Hospital for Sick Children for their advice and scientific guidance.

## Author Contributions

SW designed the experiments. SW and SE conducted all the molecular work and analyzed the data. ZT assisted in exosome fractionation experiments. MAP provided the human milk samples. MAS provided the bovine milk samples. ASF provided material support. POM provided necessary equipment and resources and supervised the research. SW and SE wrote the first draft of the paper, and all authors contributed to revising the manuscript.

## Availability of data and materials

The data used in this study are available from the corresponding author upon request.

## Supplementary Figure Captions

**Supplementary Figure 1.** Pre-processing of whole bovine milk prior to long-term storage at −80 °C. Group 1) Unprocessed whole milk stored directly at −80 °C. Group 2) Samples without cream fat globules. Group 3) Whey fractions without cream, fat globules and casein proteins.

**Supplementary Figure 2.** Bovine and human milk-derived exosome isolation via enhanced ExoQuick precipitation and ultracentrifugation methods.

**Supplementary Figure 3.** Optimized sample preparation protocol for Transmission Electron Microscopy (TEM) for the visualization of milk-derived exosomes. Copper grids were negatively stained with 2 % uranyl acetate (UA).

**Supplementary Figure 4.** Four optimized protocols, 1) QIAzol + miRNeasy MiniKit, 2) TRIzol LS, 3) TRIzol + RNA Clean and Concentrator Kit (RCC), and 4) TRIzol LS + RCC used for the isolation of total RNA from isolated milk exosomes.

