## Supplementary figures and images for "Comparison of methods for milk pre-processing, exosome isolation, and RNA extraction in bovine and human milk"

### Supplementary Fig. 1

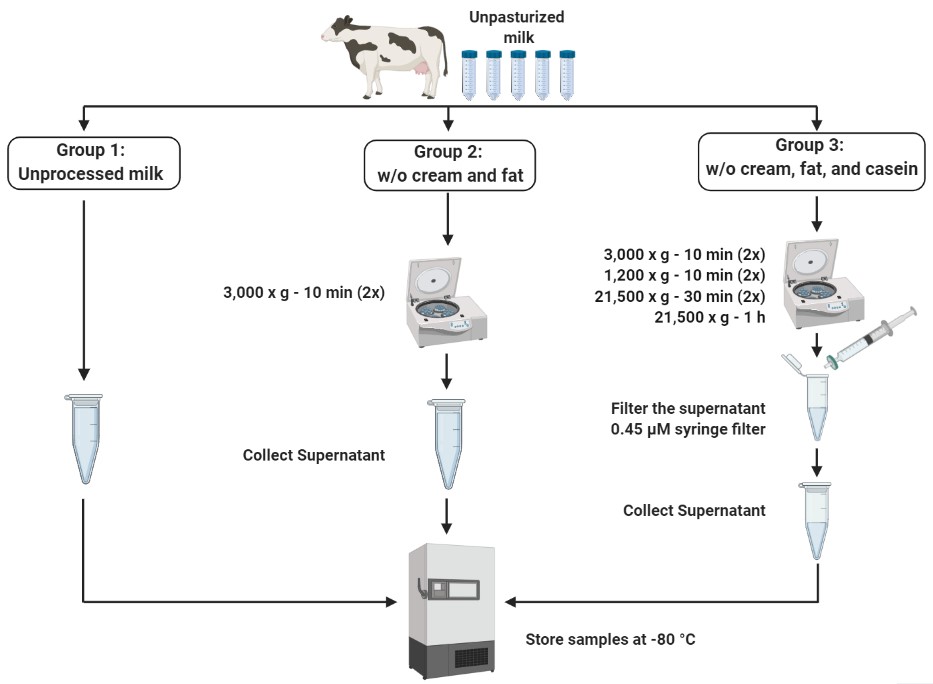

### Supplementary Fig. 2

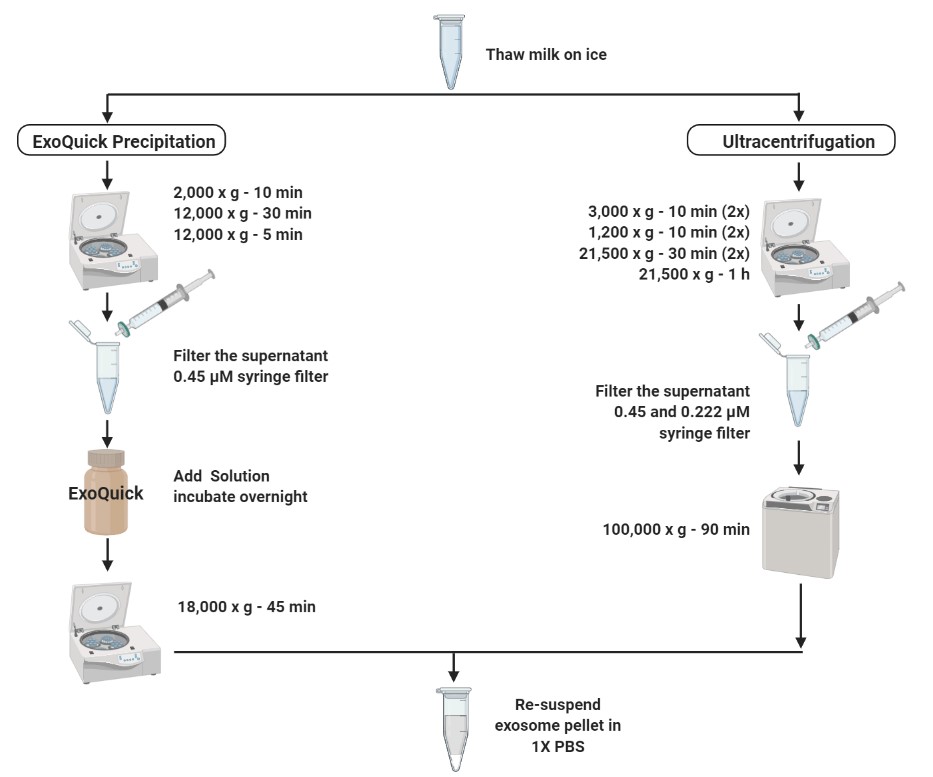

### Supplementary Fig. 3

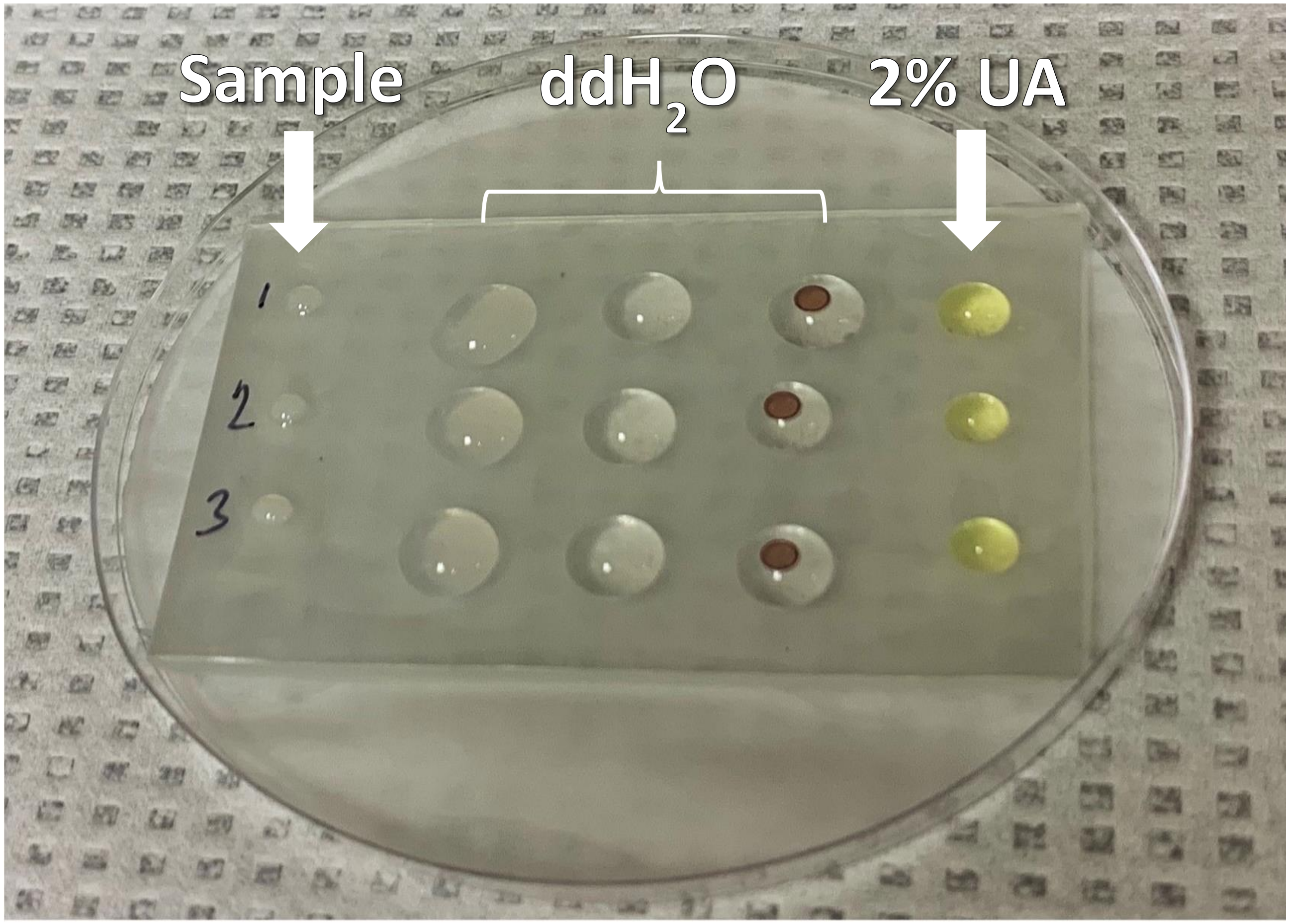

### Supplementary Fig. 4

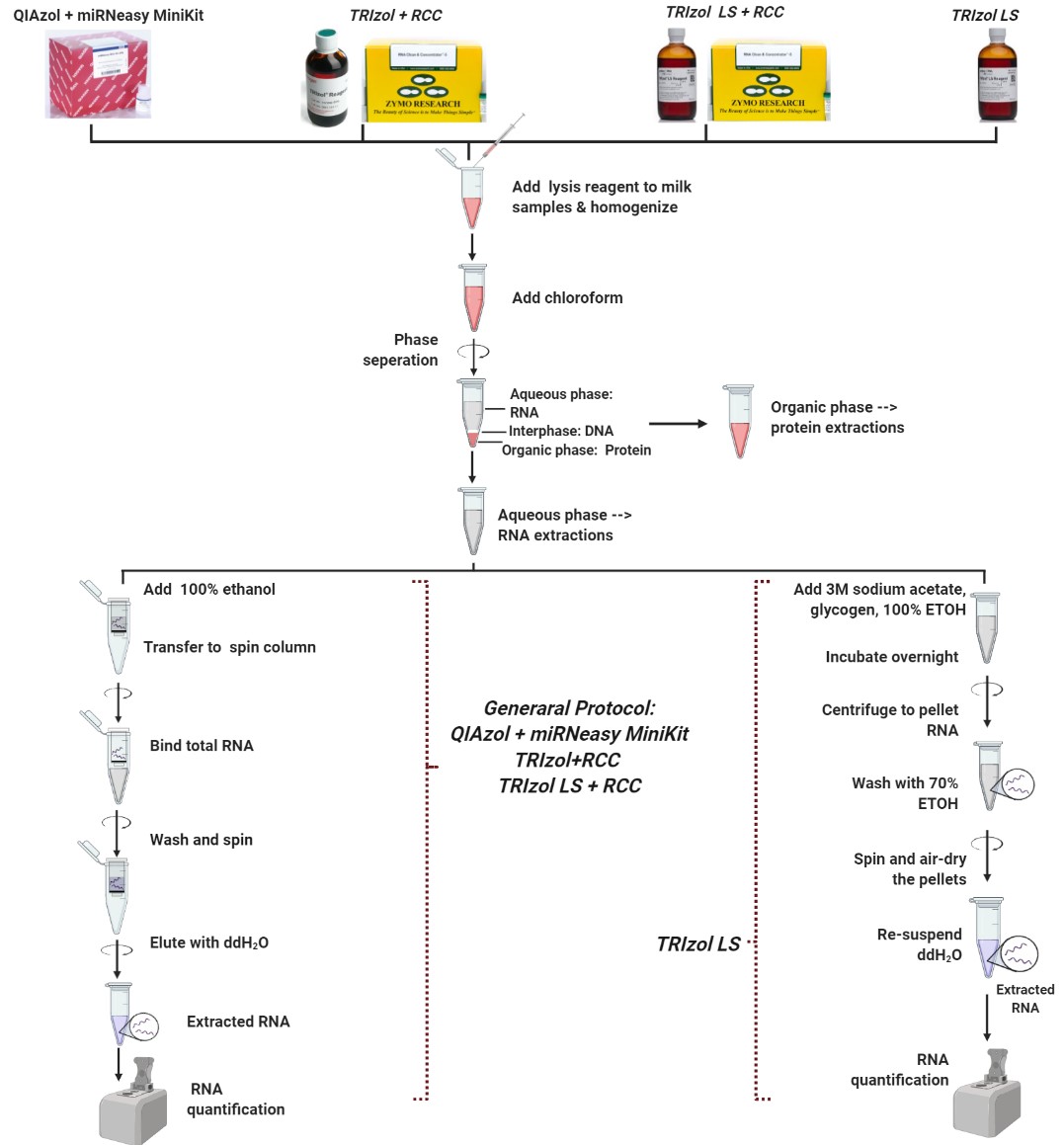
